# Guiding diffusion models for antibody sequence and structure co-design with developability properties

**DOI:** 10.1101/2023.11.22.568230

**Authors:** Amelia Villegas-Morcillo, Jana M. Weber, Marcel J.T. Reinders

## Abstract

Recent advances in deep generative methods have allowed antibody sequence and structure co-design. This study addresses the challenge of tailoring the highly variable complementarity-determining regions (CDRs) in antibodies to fulfill developability requirements. We introduce a novel approach that integrates property guidance into the antibody design process using diffusion probabilistic models. This approach allows us to simultaneously design CDRs conditioned on antigen structures while considering critical properties like solubility and folding stability. Our property-conditioned diffusion model offers versatility by accommodating diverse property constraints, presenting a promising avenue for computational antibody design in therapeutic applications. Code is available at https://github.com/amelvim/antibody-diffusion-properties.

## 1 Introduction

Antibodies are Y-shaped proteins produced by the immune system in response to pathogens called antigens [1]. Antibody engineering involves the refinement of the highly variable *complementarity-determining region* (CDR) loops to enhance its function or certain properties (Figure 1a). From a therapeutic perspective, there is a significant interest in the *in silico* design of CDRs capable of binding to specific antigens. Traditional approaches rely on energy-based optimization [2], which is computationally intensive and time-consuming. Recent advancements in deep generative methods offer enhanced performance by co-designing both the sequence and structure of CDRs simultaneously [3–5]. One notable advantage of these methods over sequence-based approaches (such as [6, 7]) is their capability to condition on both the antigen epitope and antibody framework structures during generation, which has proven useful for affinity optimization.

**Figure 1:**
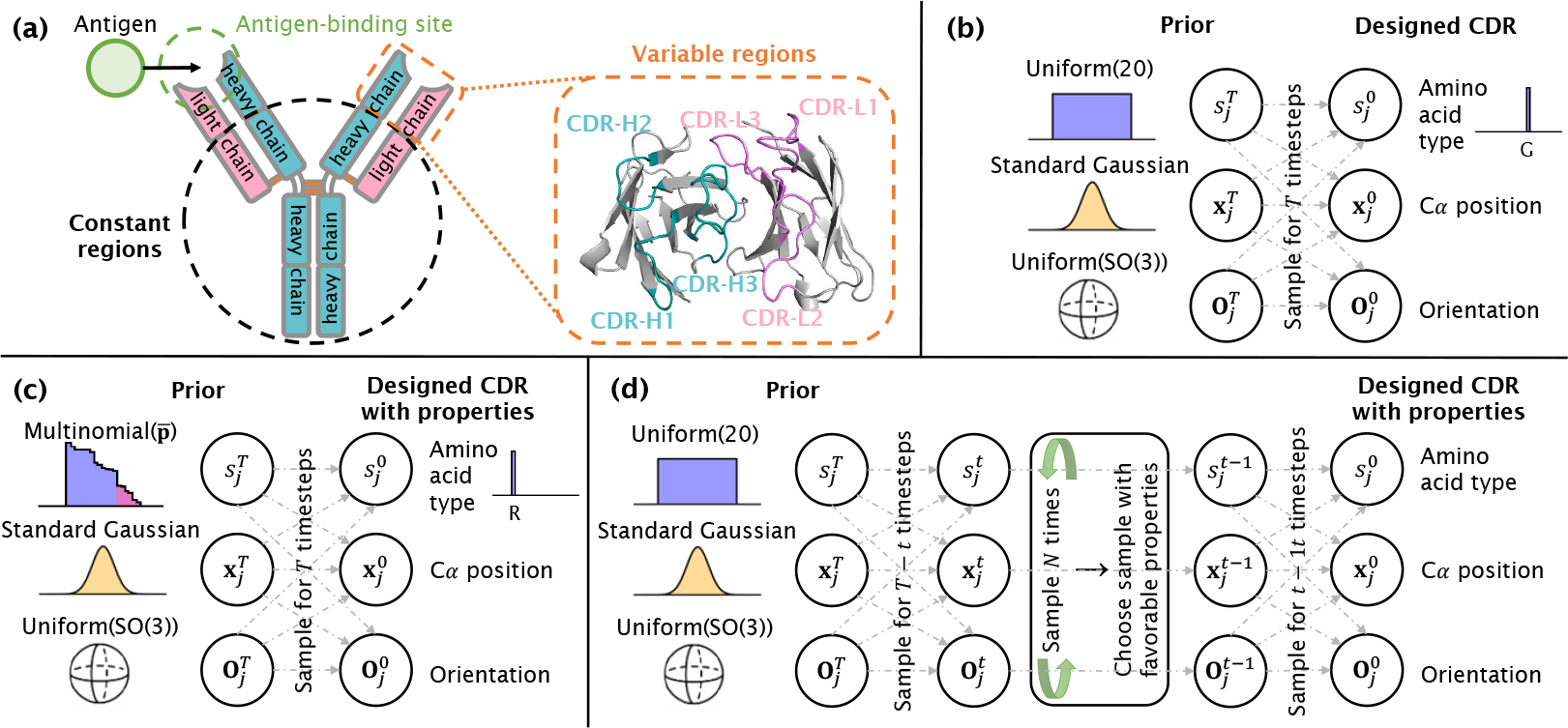
**(a)** Illustration of an antibody, featuring the two heavy (in blue) and two light (in pink) chains. The variable regions in the enlarged area encompass the antigen-binding site including the six CDR loops. **(b-d)** Visualization of the generative diffusion process, showcasing the prior distributions for each modality and the designed CDR, for the **(b)** property-unconditioned mode, **(c)** property-aware prior approach, and **(d)** sampling by property approach. (Note: The neural network parameterization is omitted from the figures but is present before sampling at each generation timestep *t*.)

Next to the antigen-targeting performance of antibodies, their *developability* properties are essential for therapeutic developments. These include factors such as solubility, aggregation propensity, thermal stability, and immunogenicity [8, 9]. These properties are vital to ensure that the antibody can be manufactured and is suitable for clinical applications [10]. While existing methods partially optimize for antigen-targeting properties, the integration of developability parameters remains an open and crucial challenge.

Consequently, in this study, we employ deep generative models for antibody design to generate *de novo* sequences and structures for the CDR loops. Beyond conditioning on the antigen structure, our approach involves guiding the model to produce candidate antibodies with favorable developability attributes. We propose a property-conditioned diffusion probabilistic model (DPM). DPMs [11, 12] have demonstrated their capability to generate realistic protein sequences and structures [13, 14], including antibodies [4]. In contrast to [15], where the diffusion process is guided with gradients from a classifier, we adopt gradient-free approaches for integrating property information. Specifically, we guide the generative diffusion process in two distinct manners: one involves incorporating a *property-aware prior*, while the other entails *sampling by property*. Notably, these approaches do not require retraining the diffusion models. Moreover, our proposed solutions are adaptable to any property or set of properties that can be computed or predicted based on the intermediate designs at the sequence or structure level. We observe that by imposing property constraints, our model yields antibodies with more favorable developability profiles while preserving their structural integrity compared to the reference antibodies.

## 2 Methods

### 2 Diffusion model for antibody design

Our work builds upon an existing method for antibody sequence and structure co-design using diffusion models. Specifically, we used the DiffAb model [4], which enables the joint generation of CDR sequences and structures while conditioning on the antibody framework and bound antigen. The model requires three inputs: amino acid types denoted as *s*_*i*_ *∈ {*ACDEFGHIKLMNPQRSTVWY*}*, C_*α*_ atom positions as **x**_*i*_ *∈* ℝ^3^, and amino acid orientations as **O**_*i*_ *∈* SO(3), where *i* is the position of the amino acid in the sequence. We generate one CDR loop at a time, denoted as *ℛ* = *{*(*s*_*j*_, **x**_*j*_, **O**_*j*_)|*j* = *l* + 1, …, *l* + *m}*, given the rest of antibody-antigen complex *C* = *{*(*s*_*i*_, **x**_*i*_, **O**_*i*_)|*i*≠ *j}*.

The forward diffusion process (*t* = 0, …, *T*) gradually introduces noise into each modality through different distributions *q* towards the prior distributions. For the amino acid types, 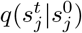 follows a multinomial distribution; for the C_*α*_ positions, 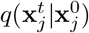 is modeled as Gaussian; and for the amino acid orientations, 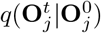 is an isotropic Gaussian. Starting from the prior distributions, the generative diffusion process (*t* = *T*, …, 0) transforms each modality toward the data distribution, as depicted in Figure 1b. In this process, parametric models *p*_*θ*_ are employed to approximate the posterior distributions at each generation timestep. Different neural networks are used for the three modalities, with a shared encoder and separate decoders. For an in-depth understanding of the diffusion process, neural network architectures, and training of the models, we refer readers to [4].

### 2.2 Antibody design guided on properties

#### Property-aware prior

The prior distribution for the amino acid types follows a uniform distribution across the 20 classes. In this approach, we propose sampling from a property-aware prior in the form:

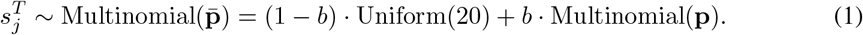

Here, **p** = [*p*_1_, …, *p*_20_], where *p*_*k*_ represents the probability of the amino acid type *k* given a property of interest. The uniform and multinomial components are weighted by a constant *b* that can be adjusted based on the application requirements. This approach is depicted in Figure 1c. The posterior probabilities at each generation timestep *t* are defined as:

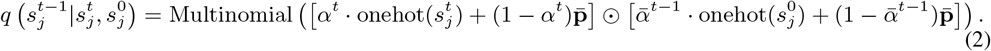

Here, 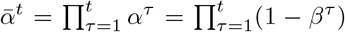 with *β*^*t*^ denoting the cosine variance schedule. 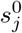 is the amino acid type approximated by the neural network model during the generative diffusion process. This posterior enforces resampling to rely more on the property-aware prior at the start of the process (*t*→ *T*), and more on the previously sampled amino acid 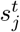 towards the end of the process (*t* → 0). Note that we need to divide the posterior probabilities by their sum to ensure that they add up to one.

Although this approach can accommodate any amino acid-related property, in this study, we focused on the hydropathy score [16] as a proxy for solubility and aggregation. Figure 4 presents this score for each amino acid type and its translation to probabilities (for different values of *b*). Here, hydrophilic amino acids (i.e. with low hydropathy scores) are assigned higher probabilities, and vice-versa.

#### Sampling by property

For properties related to sequence and structure, we developed a general guidance approach. At each generation timestep, we sample *N* times and then select the sample with the most favorable property value, as shown in Figure 1d. For instance, we choose the minimum, if our aim is to minimize the value of the property. When multiple properties are considered, we opt for the sample with the minimum sum of all property values (known as the Pareto optimal solution [17]). In a more flexible version, we convert the *N* property values into probabilities through the softmax function and then sample the next timestep from this distribution. Here, the assumption is that all *ℛ* samples generated at the same step in the process are equally valid in terms of (**s, x, O**).

For the sampling by property approach, we conditioned our model on both the hydropathy score and the folding energy (ΔG). To compute the difference in folding energy, we employed the ΔΔG predictor from [18], which relies on a graph convolutional network (GCN) model to predict the energy difference between the reference and generated antibodies. To obtain predicted ΔΔG values for the *n*-sampled CDR, we feed the model with the amino acid sequence and C_*α*_ atom positions at the current generation timestep (**s**^*t*−1^, **x**^*t*−1^), as well as those from the previous timestep (**s**^*t*^, **x**^*t*^).

### 2.3 Benchmark dataset and trained model

To benchmark our guided approaches, we employed the test set described in [4], which comprises 19 antibody-antigen complexes sourced from the SabDab database [19]. The CDR-H3 sequences of the test antibodies share a maximum of 50% sequence identity with each other and with the training data. The test set includes protein antigens from various pathogens, including influenza and SARS-CoV-2. To guide the generation diffusion process, we leveraged the `codesign_single` model from DiffAb, which has been trained to generate all CDRs, one at a time randomly selected for each training sample. Using this model, we designed single CDRs from random values provided the rest of the complex.

### 2.4 Evaluation

For each test complex, we generate 100 designs for each of the six CDRs through *T* = 100 timesteps of generation, each one maintaining the same length as the reference test CDR. We evaluate the designs using the following metrics: (i) the AAR (amino acid recovery) measures the sequence identity between the reference and generated CDR sequences, (ii) the RMSD (root mean square deviation) computes the C_*α*_ atom distance between the reference and generated CDR structures, (iii) the hydropathy score averages the hydropathy values over the generated CDR sequences, and (iv) the predicted ΔΔG [18] measures the difference in folding energy (ΔG) between the reference and generated CDRs, considering atoms (N, C_*α*_, C, O) after reconstructing the backbone structure from C_*α*_ atom positions and orientations (see [4]). We use predicted ΔΔG as it is computationally more efficient and has moderate to high correlation with experimental measures of antibody energy upon mutations [18]. For AAR, higher values are preferable, while lower values are desired for RMSD, hydropathy score, and predicted ΔΔG. Note that we aim to generate CDRs with improved property values (low hydropathy score and predicted ΔΔG) without deteriorating the structural integrity (we expect slight deviations in AAR and RMSD to the reference).

## 3 Results

### 3.1 Guidance on properties is effective

We assess our property-aware prior approach using the hydropathy score, with *b* = 0.8 (see Figure 5 for the impact of *b* on the hydropathy score of the final designs). The sampling by property approach is tested for both, the hydropathy score and ΔΔG. We select the sample with minimum ΔΔG in *N* = 20 samples at each generation timestep, as suggested by Figure 6. Additionally, we test the combination of both properties in two ways: sampling by ΔΔG with a hydropathy-aware prior, and jointly sampling by ΔΔG and hydropathy score.

Figure 2 illustrates the performance metrics for intermediate CDR-H3 designs at every 10 timesteps of generation. As observed, sampling by hydropathy results in a bigger change in the hydropathy score and the AAR compared to using a hydropathy-aware prior, even with a high value of *b*. This indicates that the generated CDR sequences differ more substantially from the reference when sampling by hydropathy. Furthermore, we note that in comparison to the unconditioned mode, exclusive sampling by ΔΔG improves the hydropathy score, whereas exclusive sampling by hydropathy does the same for the predicted ΔΔG. When both properties are combined, the most favorable outcomes are achieved, with the majority of designs exhibiting hydropathy scores and predicted ΔΔG values below zero. This is supported by Mann-Whitney statistical tests, revealing significant differences in the final metric distributions (at *t* = 0) across different models for the entire test set (see Figure 7). Meanwhile, the values of AAR and RMSD are consistent across models, which is desirable to avoid significant deviations from the reference CDR. While the sequence similarities within guided designs deviate from the unconditioned ones (as expected, Figure 8), the RMSD values remain close (Figure 9).

**Figure 2:**
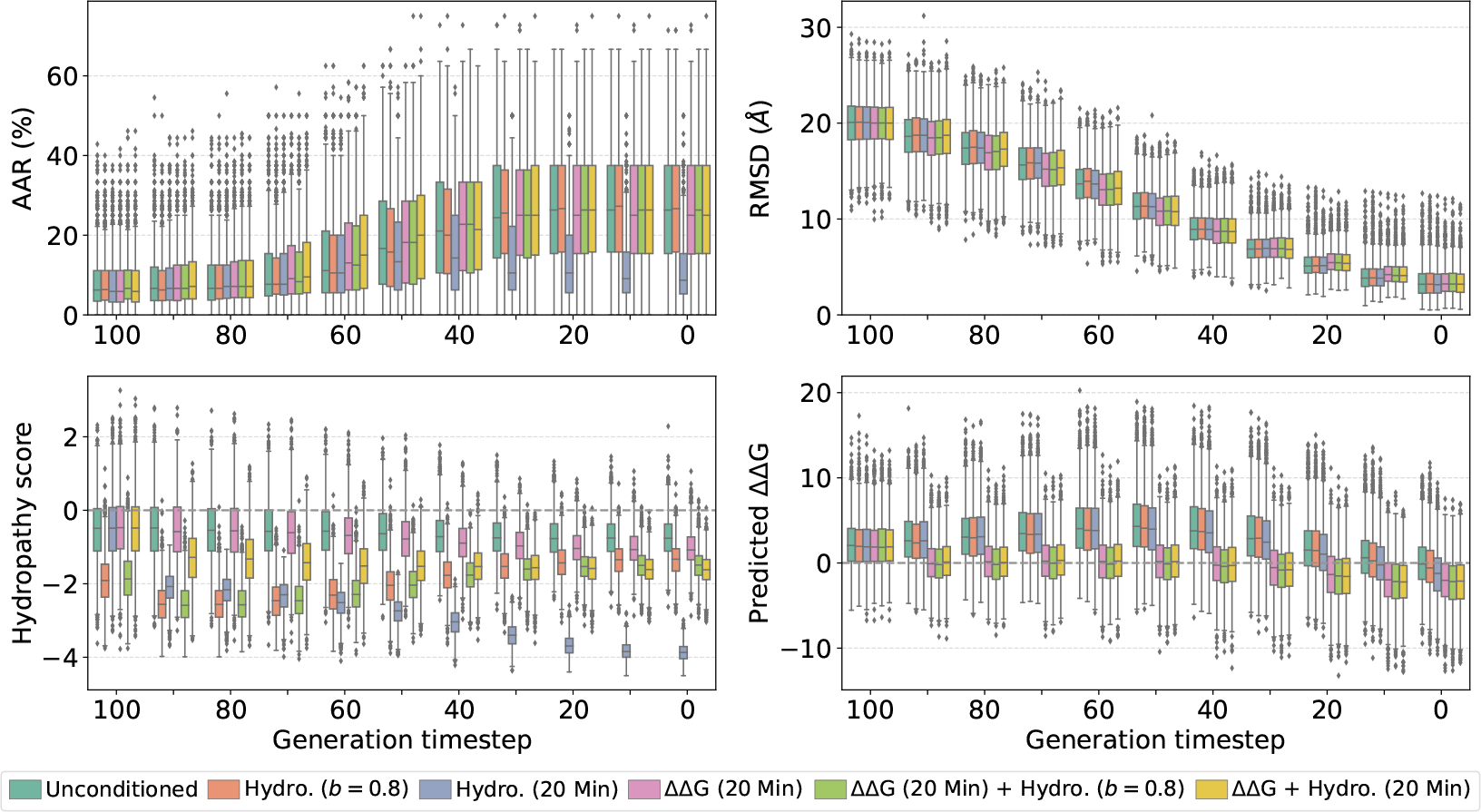
Per-timestep metrics on the 19 test complexes (design CDR-H3). The boxplots represent the distribution of metric values (AAR, RMSD, hydropathy score, and predicted ΔΔG) over 100 designed CDRs for each test complex. Here we compare the unconditioned mode with different property-guided models: hydropathy-aware prior, sampling by hydropathy or ΔΔG, and combinations of both.

These observations apply to all other CDRs as well. Figure 10 displays per-timestep metrics and Table 1 includes the performance metrics for the final designs of all CDRs. We can see that AAR decreases and RMSD increases slightly after guidance, which is expected. Furthermore, each CDR exhibits some variations in response to the guidance. For CDR-H2 and L2, sampling by hydropathy results in significantly lower AAR, leading to better predicted ΔΔG compared to sampling by ΔΔG.

**Table 1:**
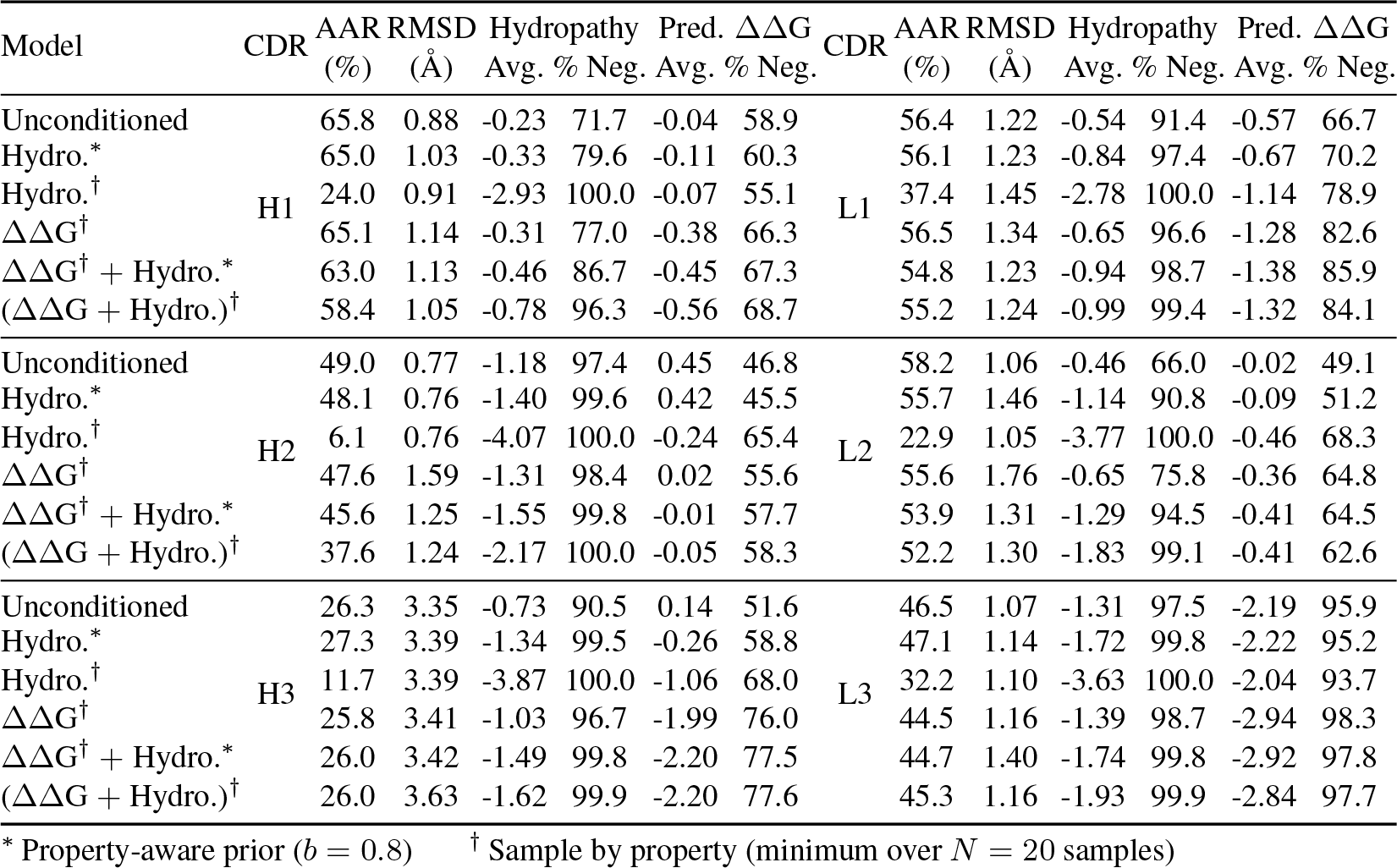
Average performance metrics over 100 designs for the 19 test complexes (for each CDR). The metrics are AAR, RMSD, hydropathy score, and predicted ΔΔG. For the hydropathy score and predicted ΔΔG, we also show the percentage of samples with negative values. Here we compare the unconditioned mode with different property-guided models: hydropathy-aware prior, sampling by hydropathy or ΔΔG, and combinations of both.

| Model | CDR | AAR (%) | RMSD (Å) | Hydropathy Avg. | Hydropathy % Neg. | Pred. $\Delta\Delta G$ Avg. | Pred. $\Delta\Delta G$ % Neg. | CDR | AAR (%) | RMSD (Å) | Hydropathy Avg. | Hydropathy % Neg. | Pred. $\Delta\Delta G$ Avg. | Pred. $\Delta\Delta G$ % Neg. |
| --- | --- | --- | --- | --- | --- | --- | --- | --- | --- | --- | --- | --- | --- | --- |
| Unconditioned | H1 | 65.8 | 0.88 | -0.23 | 71.7 | -0.04 | 58.9 | L1 | 56.4 | 1.22 | -0.54 | 91.4 | -0.57 | 66.7 |
| Hydro.* |  | 65.0 | 1.03 | -0.33 | 79.6 | -0.11 | 60.3 |  | 56.1 | 1.23 | -0.84 | 97.4 | -0.67 | 70.2 |
| Hydro.† |  | 24.0 | 0.91 | -2.93 | 100.0 | -0.07 | 55.1 |  | 37.4 | 1.45 | -2.78 | 100.0 | -1.14 | 78.9 |
| $\Delta\Delta G^\dagger$ | | 65.1 | 1.14 | -0.31 | 77.0 | -0.38 | 66.3 | | 56.5 | 1.34 | -0.65 | 96.6 | -1.28 | 82.6 |
| $\Delta\Delta G^\dagger + \text{Hydro}^*$ | | 63.0 | 1.13 | -0.46 | 86.7 | -0.45 | 67.3 | | 54.8 | 1.23 | -0.94 | 98.7 | -1.38 | 85.9 |
| $(\Delta\Delta G + \text{Hydro}.)^\dagger$ | | 58.4 | 1.05 | -0.78 | 96.3 | -0.56 | 68.7 | | 55.2 | 1.24 | -0.99 | 99.4 | -1.32 | 84.1 |
| Unconditioned | H2 | 49.0 | 0.77 | -1.18 | 97.4 | 0.45 | 46.8 | L2 | 58.2 | 1.06 | -0.46 | 66.0 | -0.02 | 49.1 |
| Hydro.* |  | 48.1 | 0.76 | -1.40 | 99.6 | 0.42 | 45.5 |  | 55.7 | 1.46 | -1.14 | 90.8 | -0.09 | 51.2 |
| Hydro.† |  | 6.1 | 0.76 | -4.07 | 100.0 | -0.24 | 65.4 |  | 22.9 | 1.05 | -3.77 | 100.0 | -0.46 | 68.3 |
| $\Delta\Delta G^\dagger$ | | 47.6 | 1.59 | -1.31 | 98.4 | 0.02 | 55.6 | | 55.6 | 1.76 | -0.65 | 75.8 | -0.36 | 64.8 |
| $\Delta\Delta G^\dagger + \text{Hydro}^*$ | | 45.6 | 1.25 | -1.55 | 99.8 | -0.01 | 57.7 | | 53.9 | 1.31 | -1.29 | 94.5 | -0.41 | 64.5 |
| $(\Delta\Delta G + \text{Hydro}.)^\dagger$ | | 37.6 | 1.24 | -2.17 | 100.0 | -0.05 | 58.3 | | 52.2 | 1.30 | -1.83 | 99.1 | -0.41 | 62.6 |
| Unconditioned | H3 | 26.3 | 3.35 | -0.73 | 90.5 | 0.14 | 51.6 | L3 | 46.5 | 1.07 | -1.31 | 97.5 | -2.19 | 95.9 |
| Hydro.* |  | 27.3 | 3.39 | -1.34 | 99.5 | -0.26 | 58.8 |  | 47.1 | 1.14 | -1.72 | 99.8 | -2.22 | 95.2 |
| Hydro.† |  | 11.7 | 3.39 | -3.87 | 100.0 | -1.06 | 68.0 |  | 32.2 | 1.10 | -3.63 | 100.0 | -2.04 | 93.7 |
| $\Delta\Delta G^\dagger$ | | 25.8 | 3.41 | -1.03 | 96.7 | -1.99 | 76.0 | | 44.5 | 1.16 | -1.39 | 98.7 | -2.94 | 98.3 |
| $\Delta\Delta G^\dagger + \text{Hydro}^*$ | | 26.0 | 3.42 | -1.49 | 99.8 | -2.20 | 77.5 | | 44.7 | 1.40 | -1.74 | 99.8 | -2.92 | 97.8 |
| $(\Delta\Delta G + \text{Hydro}.)^\dagger$ | | 26.0 | 3.63 | -1.62 | 99.9 | -2.20 | 77.6 | | 45.3 | 1.16 | -1.93 | 99.9 | -2.84 | 97.7 |

\* Property-aware prior ( $b = 0.8$ )† Sample by property (minimum over $N = 20$ samples)

### 3.2 Amino acid composition changes with hydropathy guidance

Hydropathy guidance aims to design CDR sequences containing hydrophilic amino acid types without deteriorating the target binding affinities. This effect is illustrated in the amino acid compositions in Figure 11. We note that sampling by hydropathy causes the most significant shift in the final amino acid distribution toward arginine (R) and aspartic acid (D), two of the most hydrophilic amino acids. Furthermore, this approach eliminates most of the hydrophobic amino acids. Using the hydropathy-aware prior, the effect is not as strong, primarily because the model relies less on the prior towards the end of the generation process. Sampling by ΔΔG, whether alone or in combination with hydropathy, also increases the number of hydrophilic amino acids, such as tyrosine (Y). These results, along with Table 1, indicate a correlation between both properties. Improved hydropathy profiles lead to larger exposed surface areas, which are necessary for the antibody-antigen interaction.

### 3.3 Energy distributions shift with ΔΔG guidance, even after relaxation

The objective of ΔΔG guidance is to generate CDR loops with enhanced folding stability, leading to potential improvement in antibody-antigen binding. Figure 3a shows the relationship between predicted ΔΔG and hydropathy score for the final CDR-H3 designs, revealing the positive correlation between these two properties. Considering that lower values are desirable for both properties, we calculated the Pareto frontiers for the three approaches. Notably, we observe that the three frontiers are clearly separated, with the guided approaches exhibiting a trend towards the lowest values. Thus, they outperform a naive filter on top-scoring samples from the unconditioned model. The most favorable Pareto solutions are obtained when jointly sampling by ΔΔG and hydropathy. The empirical run-time comparisons for this test complex are in Table 2.

**Table 2:** Empirical run-time comparisons for test complex 7chf_A_B_R (design CDR-H3). While modifications to the prior incur no time overhead, the speed of the sampling approach depends on the predictor. Hydropathy computation is fast, but the ΔΔG predictor has a much slower execution time.

| Model | Full (100 designs, 100 timesteps) | One batch (16 designs), one timestep |  |
| --- | --- | --- | --- |
|  |  | Get predictions | Denoise (and sample<sup>†</sup>) |
| Unconditioned and Hydro.* | 360.21 sec (6 min) | 0.6304 sec | 0.0063 sec |
| Hydro.<sup>†</sup> | 436.29 sec (7.3 min) |  | 0.1248 sec |
| $\Delta\Delta G^\dagger$ and $\Delta\Delta G^\dagger + \text{Hydro.}^*$ | 3275.72 sec (54.6 min) | | 4.5499 sec |
| $(\Delta\Delta G + \text{Hydro.})^\dagger$ | 3280.60 sec (54.7 min) | | 4.6691 sec |

\* Property-aware prior ( $b = 0.8$ )
<sup>†</sup> Sample by property (minimum over $N = 20$ samples)

**Figure 3:**
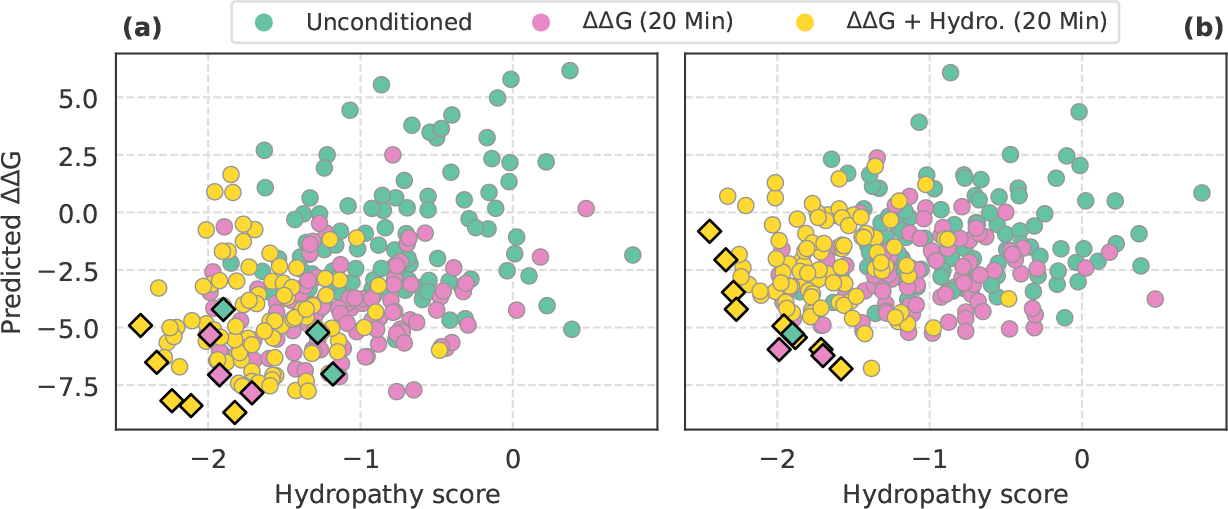
Distribution of hydropathy scores and predicted ΔΔG for test complex 7chf_A_B_R (design CDR-H3), **(a)** before and **(b)** after Rosetta relaxation. The highlighted points (diamond markers) correspond to the Pareto optimal solutions.

**Figure 4:**
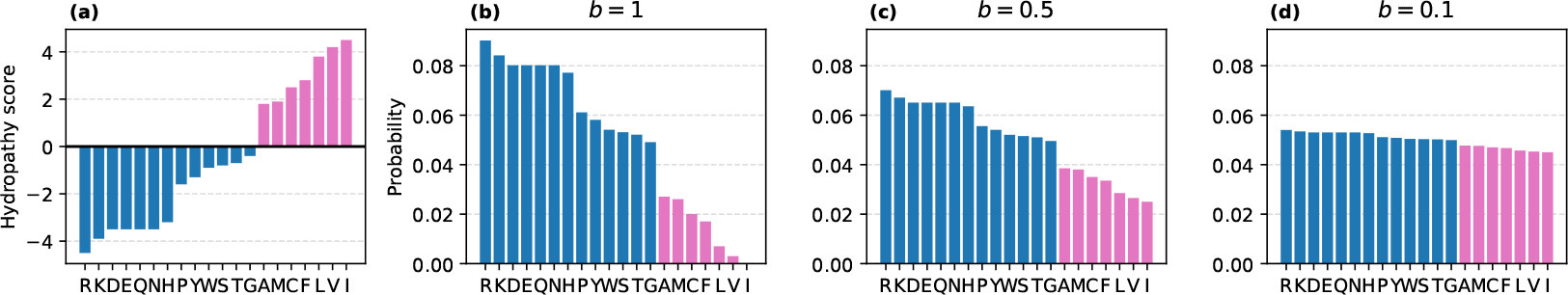
**(a)** Hydropathy scores per amino acid and conversion to probability distribution. **(b-d)** Combined uniform-hydropathy distribution for different values of *b*. Amino acid types are ordered by ascending hydropathy score.

**Figure 5:**
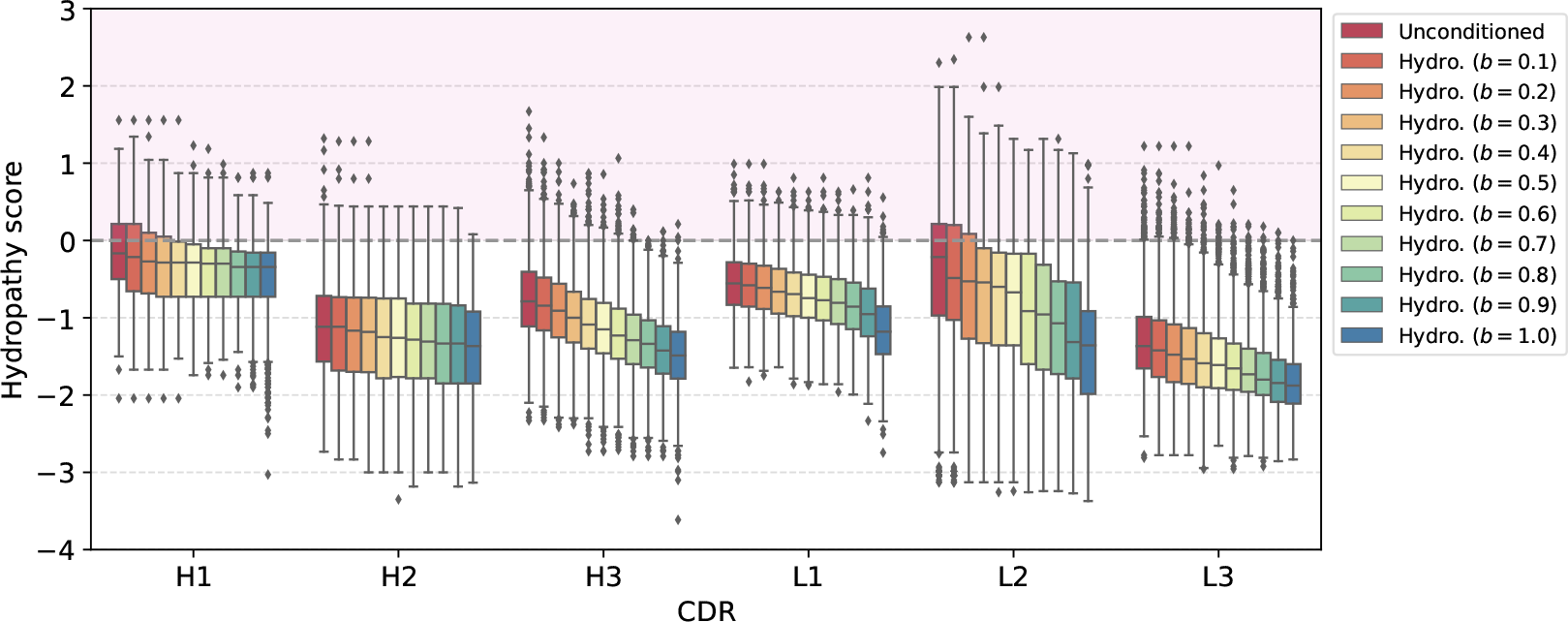
Distribution of hydropathy scores over 100 designed CDRs for each test complex (total 19) when conditioning on hydropathy prior with different values of *b*. We observe that the unconditioned model already generates samples with a negative median hydropathy. However, adding this prior results in a shift of the sample distributions towards more negative values of hydropathy (i.e. more hydrophilic amino acids), particularly for CDRs H3, L1, L2, and L3.

**Figure 6:**
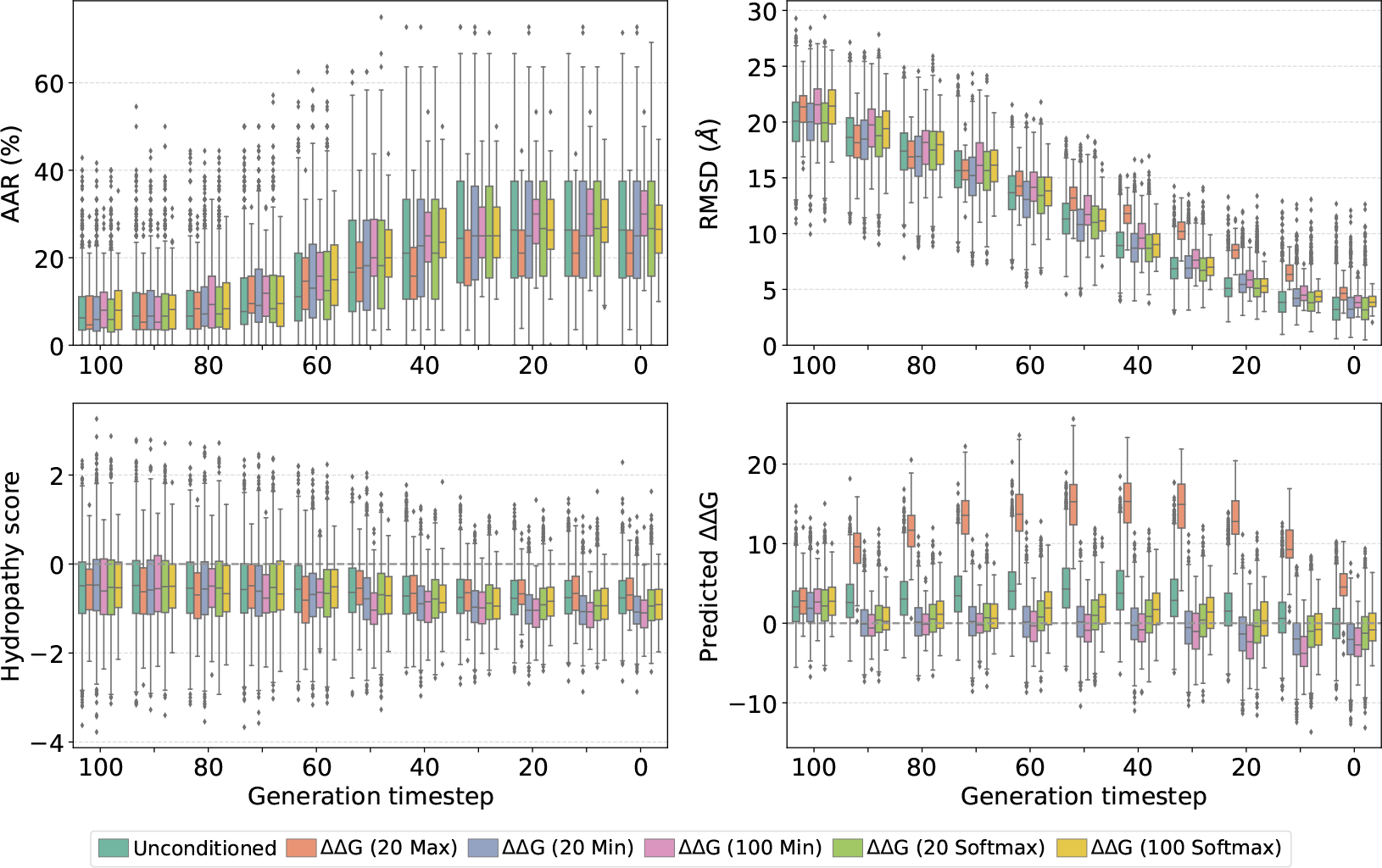
Per-timestep metrics for test complex 5xku_C_B_A (design CDR-H3). The boxplots represent the distribution of metric values (AAR, RMSD, hydropathy score, and predicted ΔΔG) over 100 designed CDRs. Here we compare different options for sampling by ΔΔG, as the number of samples *N* to obtain at every timestep, and the selection criteria (min., max., or softmax). As we can see, the max. option has a negative effect on all metrics (particularly RMSD and predicted ΔΔG), whereas taking the min. over *N* = 20 samples improves the hydropathy score and predicted ΔΔG, without affecting the values of RMSD and AAR. Instead, using *N* = 100 samples, as well as softmax-sampling are options that do not seem to help overall.

**Figure 7:**
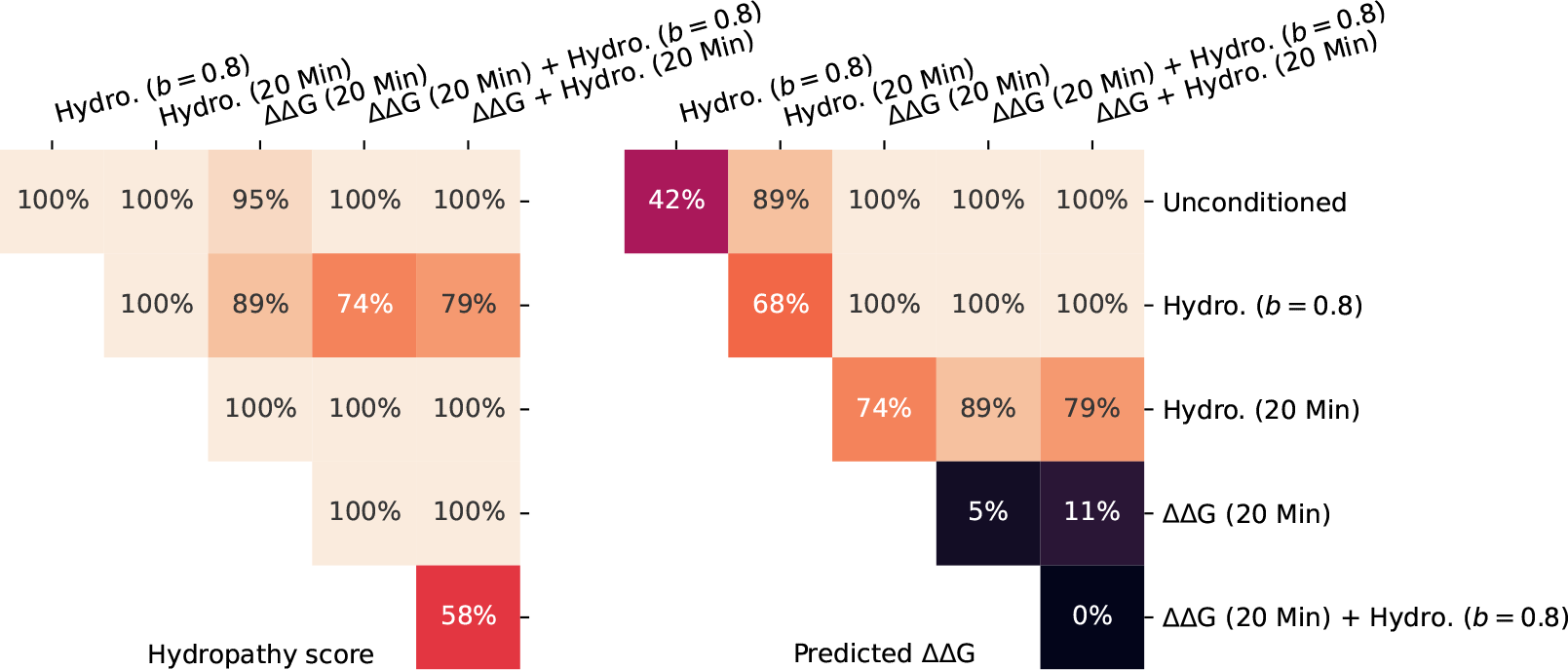
Percentage of test complexes (total 19, design CDR-H3) with significantly different metric distributions (hydropathy score or predicted ΔΔG) determined using the Mann-Whitney statistical test for every pair of models (*p*-value *<* 0.05, Benjamini-Hochberg). For most test complexes, the distributions of metrics from guided models are statistically different from those of the unconditioned mode. One exception is the two combinations of properties and the predicted ΔΔG metric.

**Figure 8:**
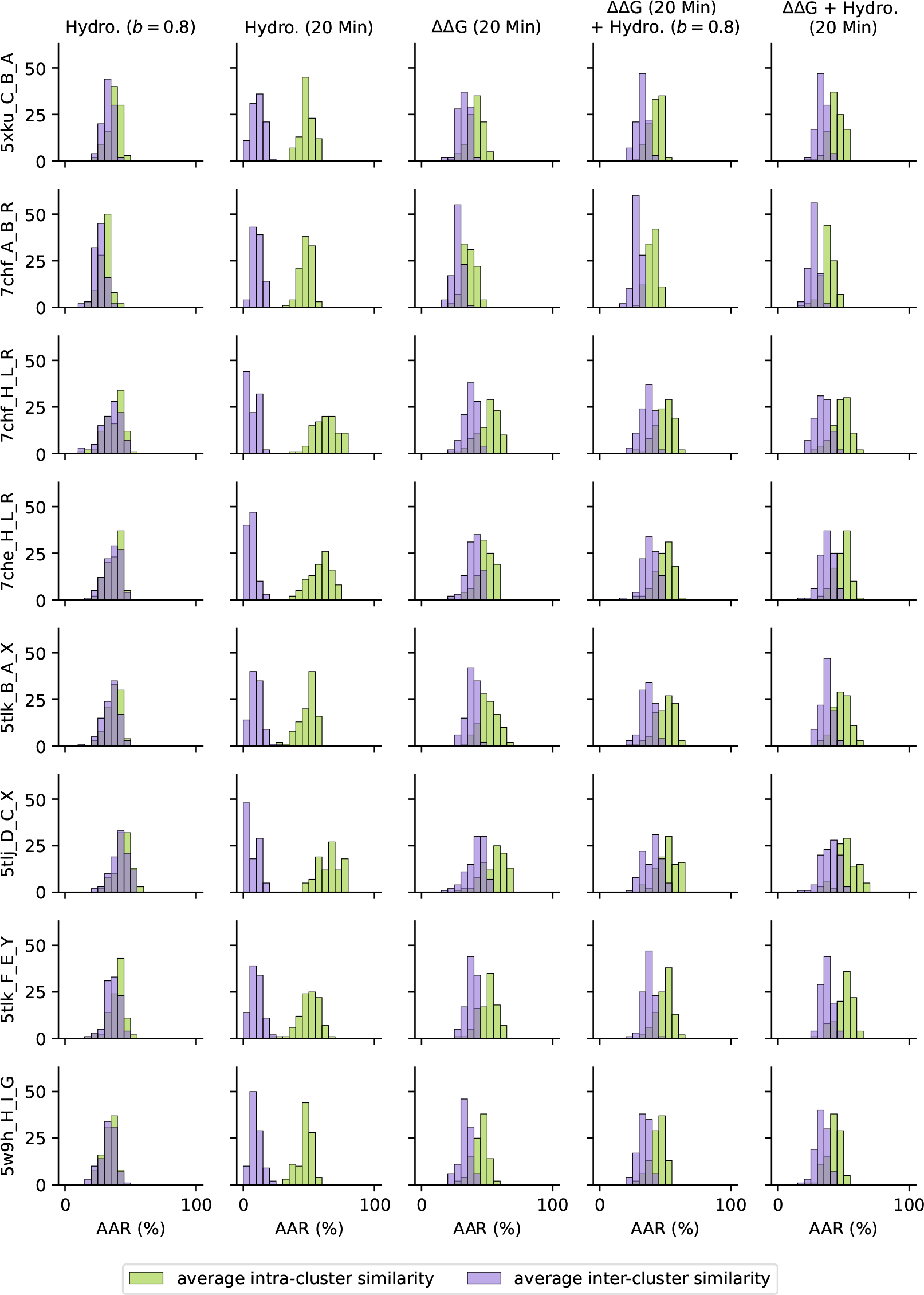
Average intra- and inter-cluster sequence similarity (measured by amino acid recovery, AAR) among the 100 CDR-H3 designs per test complex, comparing each of the guided approaches to the unconditioned mode. For most guided approaches, we observe that intra-cluster similarities are higher than inter-cluster similarities. This indicates that the conditionally-designed CDR sequences cluster closely with the unconditioned designs, but do not fully overlap. Two exceptions are the hydropathy-aware prior and sampling by hydropathy, where both clusters either overlap or are fully separated, respectively.

**Figure 9:**
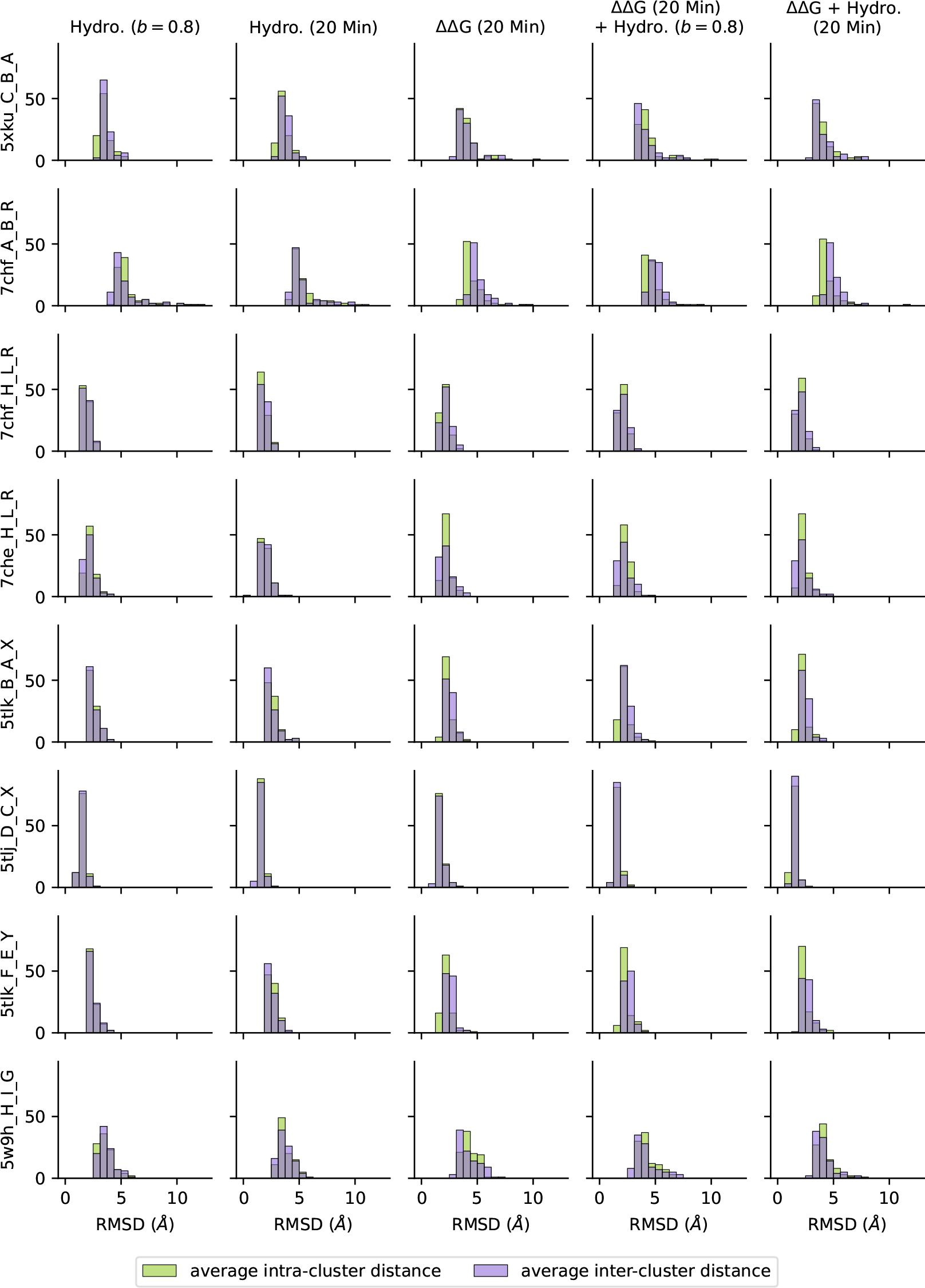
Average intra- and inter-cluster structure distance (measured by root mean square deviation, RMSD) among the 100 CDR-H3 designs per test complex, comparing each of the guided approaches to the unconditioned mode. For all guided approaches, we observe highly similar intra- and inter-cluster distances, indicating that the conditionally-designed CDR structures cluster together with the unconditioned designs.

**Figure 10:**
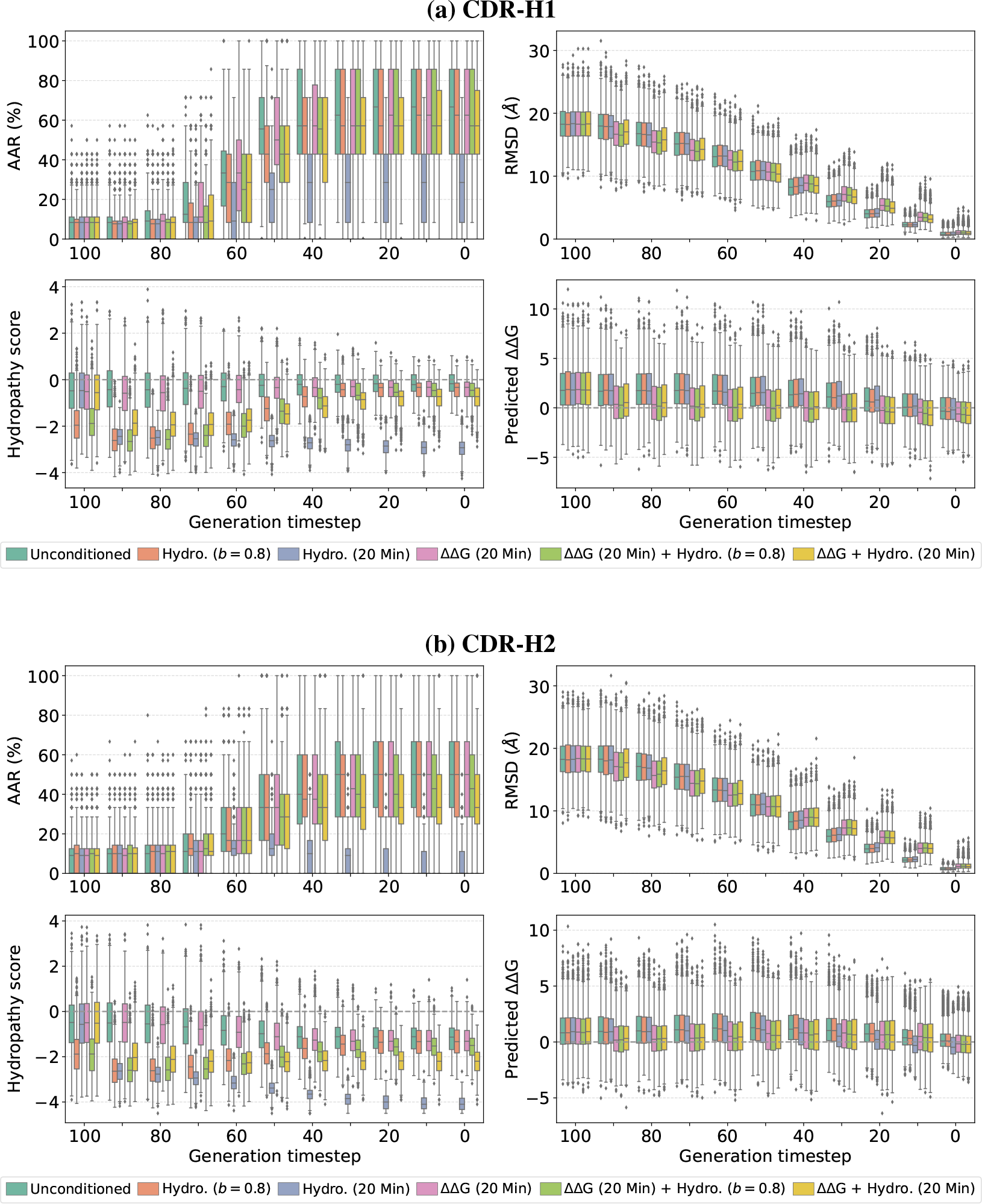
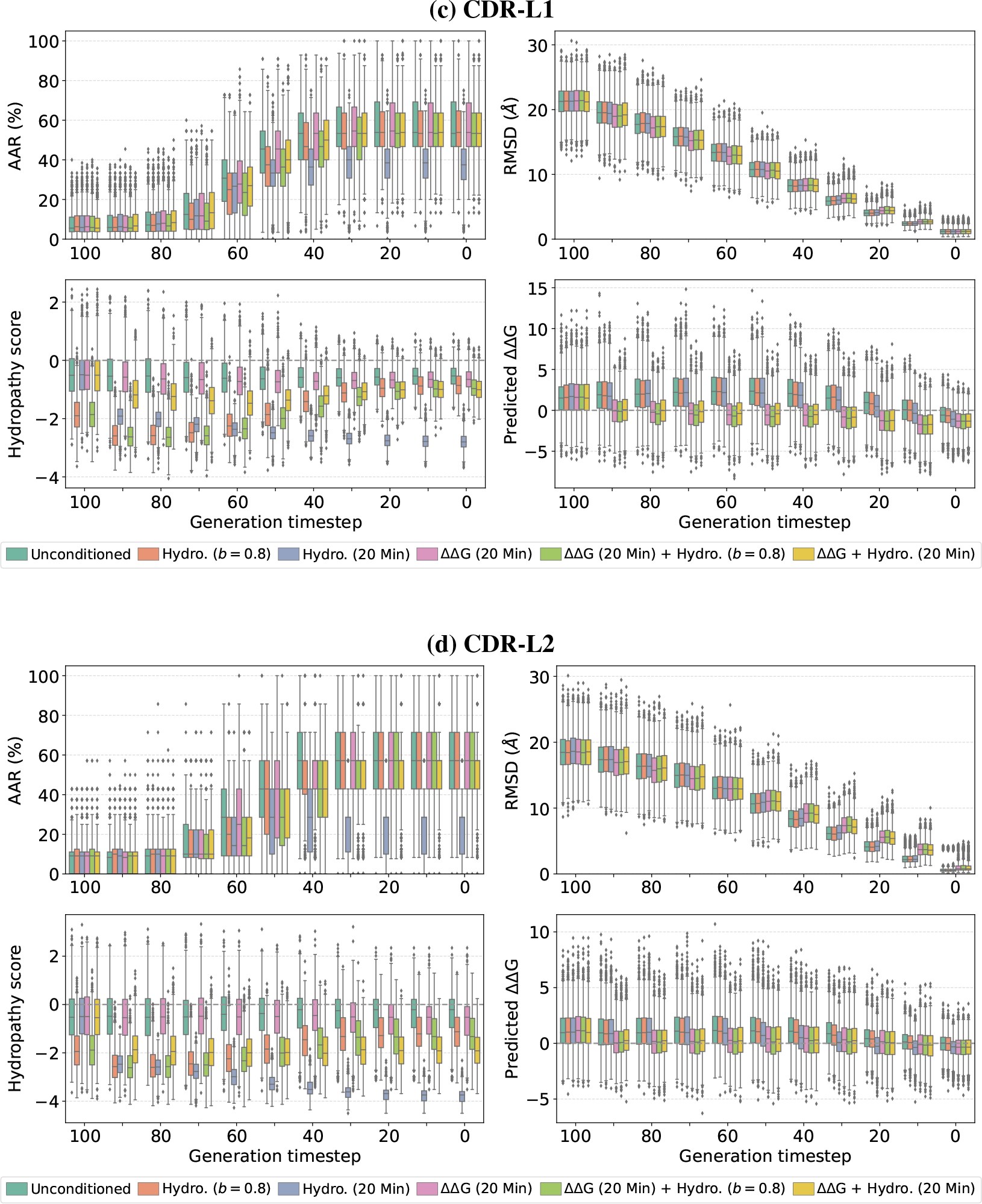
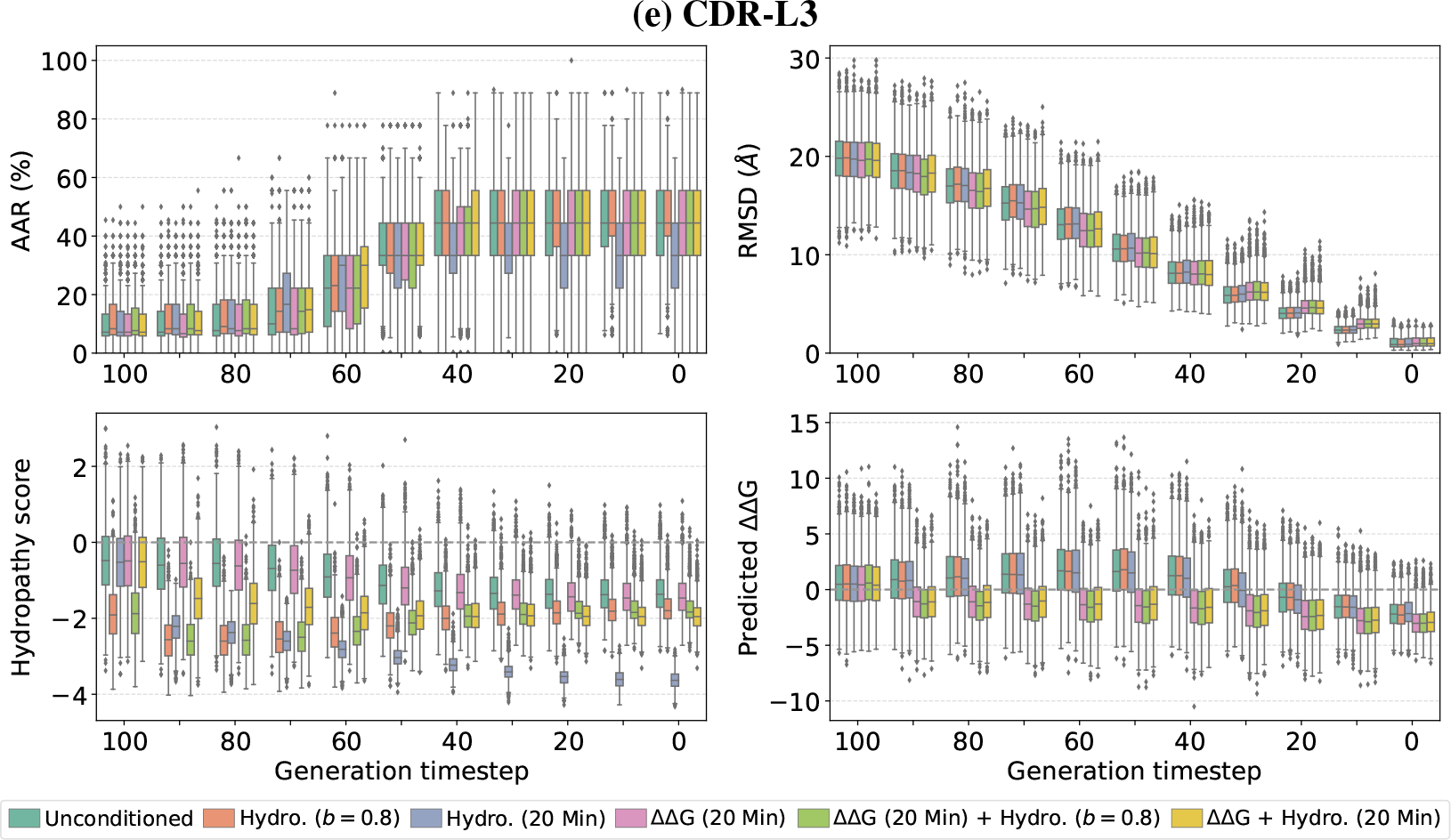
Per-timestep metrics on the 19 test complexes (design **(a)** CDR-H1 and **(b)** CDR-H2). The boxplots represent the distribution of metric values (AAR, RMSD, hydropathy score, and predicted ΔΔG) over 100 designed CDRs for each test complex. Here we compare the unconditioned mode with different property-guided models: hydropathy-aware prior, sampling by hydropathy or ΔΔG, and combinations of both. Per-timestep metrics on the 19 test complexes (design **(c)** CDR-L1 and **(d)** CDR-L2). Per-timestep metrics on the 19 test complexes (design **(e)** CDR-L3).

**Figure 11:**
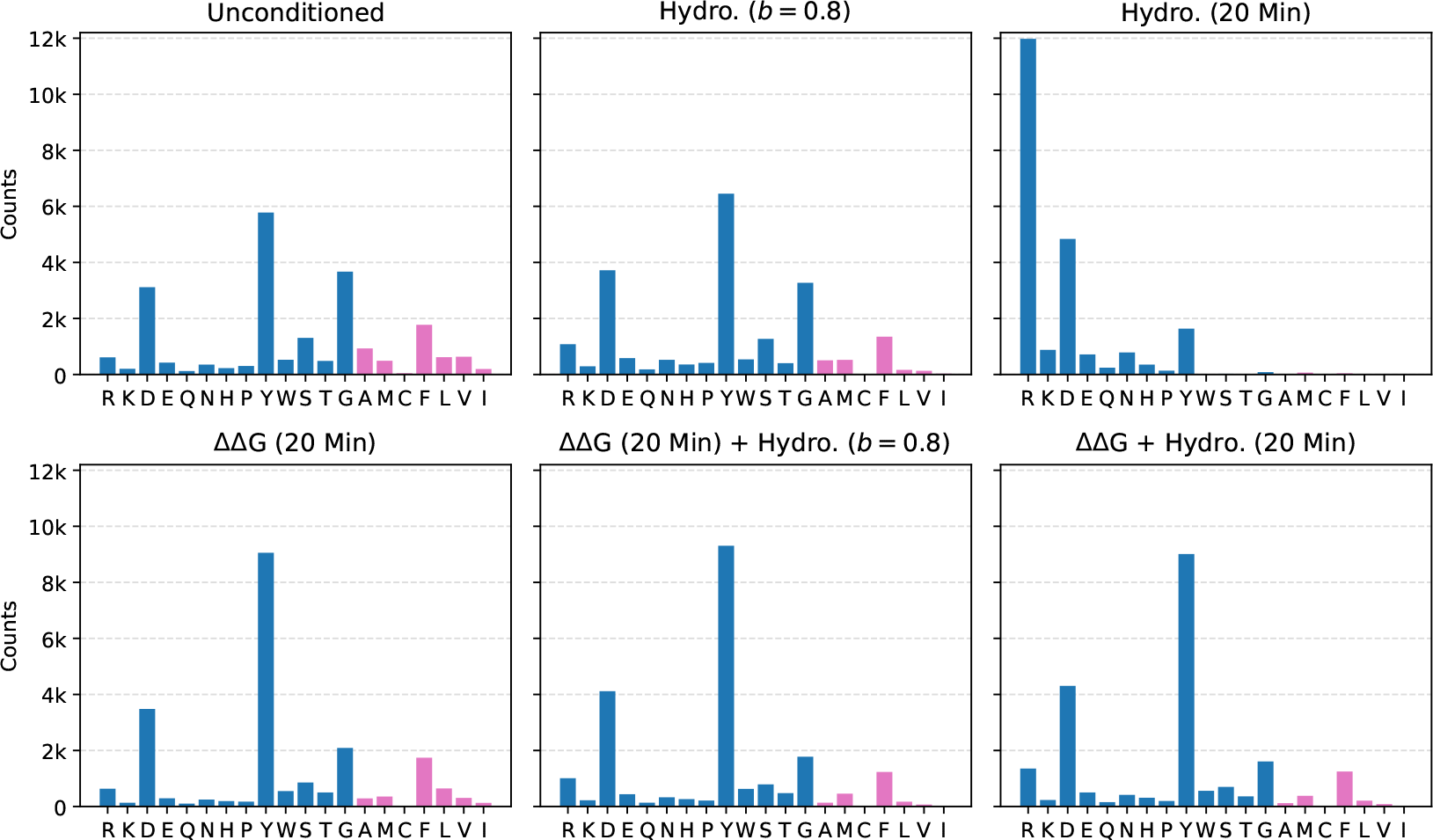
Amino acid composition for the 19 test complexes (100 CDR-H3 designs each). Amino acid types are ordered by ascending hydropathy score, while counts are colored by negative (blue) or positive (pink) hydropathy.

We then reconstruct the side-chain atoms using Rosetta [20], resulting in a refined structure. Figure 3b shows that even though the three Pareto frontiers become closer after relaxation, the distributions of the guided models are still nearer to the lower-left part than the unconditioned mode. Furthermore, compared to pre-relaxation, we attain a larger number of Pareto optimal solutions for the combined sampling by ΔΔG and hydropathy. The performance metrics for both the pre- and post-relaxed designs are in Table 3. For this example, we also visualize the resulting CDR-H3 structures in relation to the antigen epitope. We select those designs that are present in both Pareto frontiers, before and after relaxation. As observed in Figure 12, different CDR sequences lead to similar structures compared to the reference, but exhibiting improved hydropathy and predicted ΔΔG values.

**Table 3:** Average performance metrics over 100 designs for test complex 7chf_A_B_R (design CDR-H3). The metrics are AAR, RMSD, hydropathy score, and predicted ΔΔG. For the hydropathy score and predicted ΔΔG, we also show the percentage of samples with negative values. Here we compare the unconditioned mode with two property-guided models (sampling by ΔΔG, and combined sampling by ΔΔG and hydropathy), before and after Rosetta relaxation.

| Model | AAR (%) | RMSD (Å) | Hydropathy | | Pred. $\Delta\Delta G$ | |
| --- | --- | --- | --- | --- | --- | --- |
|  |  |  | Avg. | % Neg. | Avg. | % Neg. |
| Unconditioned | 22.7 | 5.11 | -0.77 | 92.0 | -0.99 | 67.0 |
| Unconditioned – Relaxed |  | 5.18 |  |  | -1.05 | 72.0 |
| $\Delta\Delta G^\dagger$ | 21.7 | 5.04 | -1.07 | 97.0 | -4.23 | 98.0 |
| $\Delta\Delta G^\dagger$ – Relaxed | | 4.85 | | | -2.72 | 94.0 |
| $(\Delta\Delta G + \text{Hydro.})^\dagger$ | 24.3 | 5.03 | -1.67 | 100.0 | -4.70 | 97.0 |
| $(\Delta\Delta G + \text{Hydro.})^\dagger$ – Relaxed | | 4.88 | | | -2.32 | 90.0 |

<sup>†</sup> Sample by property (minimum over $N = 20$ samples)

**Figure 12:**
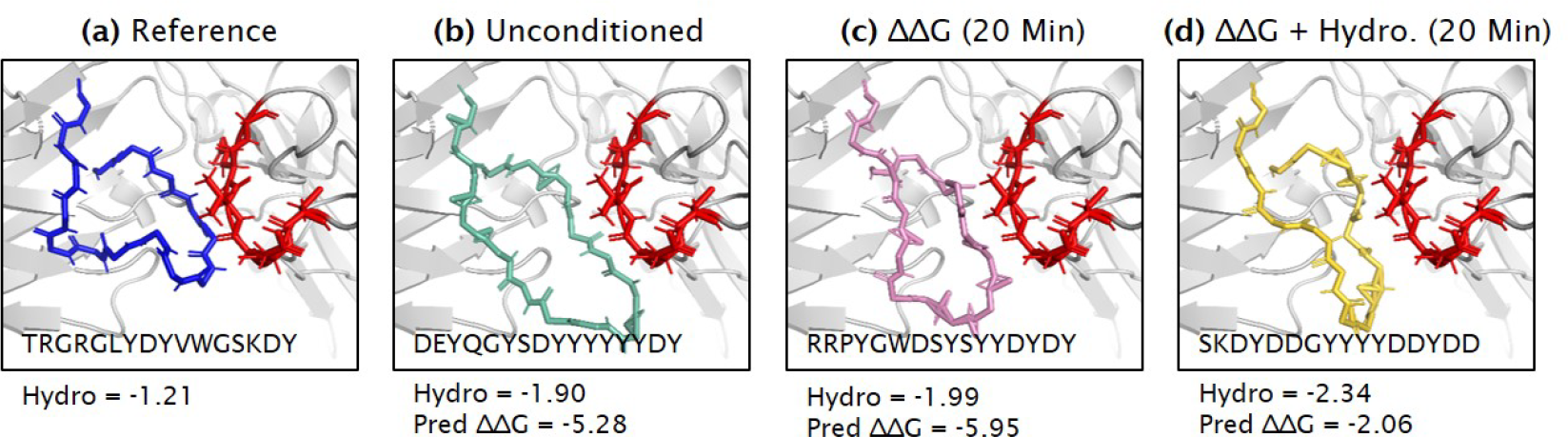
Sequence-structure designs from the Pareto frontier over hydropathy score and predicted ΔΔG after Rosetta relaxation (test complex 7chf_A_B_R, design CDR-H3). The antigen epitope is displayed in red color.

## 4 Discussion

We successfully developed two methodologically distinct strategies for conditioning diffusion models in the field of antibody design. Most notably, we can guide the generative process toward novel CDR designs with desired properties. An advantage of our approaches is their pure integration into the generative diffusion process, eliminating the need for retraining the models. While we assess our approaches using two specific properties, hydropathy and folding energy, our methodological framework can seamlessly accommodate any desired property derived from either the amino acid sequence, the structure, or both. We also demonstrate that our guided approaches enable the optimization of multiple properties at once, leading to a better set of Pareto optimal solutions. Empirical results support our modeling choices, and exploring a mathematical foundation remains of interest to better understand the validity and biases in the designs introduced by our sampling approaches. Moreover, it is crucial to note that the guided designs require experimental validation in a wet lab to confirm enhancements in solubility, aggregation, and folding/binding energy values.

## A Effect of *b* in hydropathy prior

The hydropathy scale has been proposed in [16] to take into consideration the hydrophilic (affinity to water) and hydrophobic (repelled by water) properties of each of the 20 amino acid side-chains. The hydropathy score typically ranges from −2 to +2 for most proteins, where positive values indicate higher hydrophobicity and negative values indicate higher hydrophilicity.

## B Options for sampling by ΔΔG

## C Extended results

## D Analysis of generated structures

We analyze the “designability” of the CDR-H3 generated structures for the unconditioned mode and the two guided models in Figure 12. To do so, we predict 3D structures from the anti-body sequences with designed CDR-H3 using ABodyBuilder2 (https://opig.stats.ox.ac.uk/webapps/sabdab-sabpred/sabpred/abodybuilder2/) and tFold-Ab (https://drug.ai.tencent.com/en). Table 4 contains the self-consistency RMSD (scRMSD) between the generated CDR-H3 structures (from the diffusion models) and the predicted ones from the amino acid sequences. Given that these methods have an inherent prediction error of approximately 3 Å for the CDR-H3, it is challenging to assert the accuracy of the predictions. Nonetheless, we note that the scRMSD values of the structures predicted by tFold-Ab are below 3 Å, which aligns closely with the inherent prediction error. This suggests that the generated CDR-H3 samples are likely to be “designable”.

**Table 4:** Self-consistency RMSD of designs from the Pareto frontier (test complex 7chf_A_B_R, design CDR-H3) given by predicted structures using ABodyBuilder2 and tFold-Ab. Here we compare the unconditioned mode with two property-guided models (sampling by ΔΔG, and combined sampling by ΔΔG and hydropathy).

| Model | scRMSD [ABodyBuilder2] | scRMSD [tFold-Ab] |
| --- | --- | --- |
| Unconditioned | 3.591 Å | 2.465 Å |
| $\Delta\Delta G^\dagger$ | 5.972 Å | 2.755 Å |
| $(\Delta\Delta G + \text{Hydro.})^\dagger$ | 4.536 Å | 2.578 Å |

<sup>†</sup> Sample by property (minimum over $N = 20$ samples)

